# Nisin penetrates *Staphylococcus aureus* biofilms but shows differences in killing effects against sessile and planktonic cells

**DOI:** 10.1101/303636

**Authors:** Fernanda Godoy-Santos, Betsey Pitts, Philip S. Stewart, Hilário Cuquetto Mantovani

## Abstract

Biofilms may restrict antimicrobial penetration and contribute to the recalcitrance of bacterial infections. In this work, we investigated the penetration of nisin into *S. aureus* biofilms and compared the susceptibility of *S. aureus* planktonic and sessile cells to this lantibiotic. Biofilms were grown under continuous flow in CDC reactors and calcein fluorescence was used to monitor the effect of nisin on the cytoplasmic membrane of *S. aureus* cells. Confocal scanning laser microscopy (CLSM) showed that calcein was lost within approximately 20 min in CDC biofilms, demonstrating that nisin penetrated to the bottom of the biofilm and caused membrane permeabilization. Viability analysis using PI staining showed that nisin was bactericidal against *S. aureus* sessile cells. Time-kill assays were performed against *S. aureus* in the following conditions: homogenized exponential planktonic (HEP), homogenized stationary planktonic (HSP), homogenized CDC biofilm (HB) and intact CDC biofilm (IB). The mean viability reduction of HEP and HSP were 6.71 and 1.64 log CFU.ml^-1^, respectively, confirming that stationary *S. aureus* cells were much less susceptible than exponential cells. The HB and IB treatments showed mean viability reductions of 1.25 and 0.50 log CFU.ml^-1^, respectively. Nisin activity against *S. aureus* was not limited by its ability to penetrate the bacterial biofilm, but the killing efficacy of the antimicrobial peptide was reduced by the physiological status of the biofilm-grown cells.

**Importance:** Biofilms represent a major problem to control microorganisms in industrial environments and medical devices. We developed a direct real-time microscopic visualization technique to demonstrate experimentally that the antimicrobial peptide nisin is able to penetrate *S. aureus* biofilms. Our results confirmed that nisin caused membrane permeabilization of sessile bacteria and revealed qualitative agreement between viability loss and membrane integrity loss. This approach could improve the evaluation of antibacterial susceptibility breakpoints when testing the efficacy of standard and novel antimicrobials.

## Introduction

*S. aureus* is a contagious opportunistic pathogen often associated with human and animal diseases such as mastitis, meningitis, skin and bloodstream infections (1–3). Staphylococcal infections are often characterized by a low rate of cure, high incidence, and chronicity that contributes to its high impact on the public health and worldwide economy (4, 5). The chronicity and recalcitrance of staphylococcal infections have been related to the ability of *S. aureus* to form biofilms, and previous studies demonstrated that biofilm-associated catheter infections contribute to the increasing problem of nosocomial infections (6–8). This complex mode of growth is an important virulence determinant during the establishment and maintenance of persistent infections and also confers resilience to antibiotic therapy (5, 6, 9–11).

In biofilms, bacterial cells are surrounded by an extracellular matrix composed mainly of polysaccharides, proteins, nucleic acids, and lipids. This matrix accounts for up to 90% of the dry mass of the biofilm and provides heterogeneous environments harboring microbial communities with great physiological diversity (12, 13). In addition to the presence of antimicrobial resistant and tolerant cells, the extracellular matrix may contribute to the low efficiency of strategies aiming to control established biofilms (11). Nonspecific interactions between biofilm matrix components and antimicrobials pose the potential to retard diffusion of antimicrobial agents through the biofilm (14). These events contribute to a heterogeneous distribution of inhibitory compounds within the biofilm, causing microbial cells to be exposed to sublethal doses of the antimicrobial agents (15–17). Since ineffective treatments could promote the selection of antibiotic-resistant cells and increase the incidence and persistence of bacterial infections (10, 18, 19), alternative therapeutic strategies have been developed (20, 21).

Among these strategies, bacteriocins have been proposed as a potentially useful alternative to control clinical pathogens, such as methicillin-resistant *S. aureus* (MRSA), *Enterococcus faecalis, Listeria monocytogenes*, and *Clostridium difficile* (22–24). The lantibiotics, a subgroup of bacteriocins characterized by the presence of post-translationally modified amino acid residues (ex.: lanthionine and methyllanthionine amino acid residues), has been extensively explored in that context. Their broad spectrum of activity, low cytotoxicity against mammalian cells and multiple modes of action are useful properties of lantibiotics for therapeutic applications (25–27).

Nisin is a ribosomally synthesized antimicrobial peptide that shows inhibitory activity at the nanomolar range (28), is Generally Recognized as Safe (GRAS) and is used in more than 50 countries as a biopreservative for food products (29). Nisin contains modified amino acid residues including four methyllanthionines, one lanthionine, two dehydroalanines and one dehydrobutyrine (30) that play an important role in its antimicrobial activity and receptor specificity (31–33). Nisin can bind lipid II with high affinity to form pores in the cytoplasmic membrane of target cells and also block peptidoglycan biosynthesis, thus inhibiting cell wall formation (34).

Previous studies demonstrated that nisin inhibits both planktonic and sessile cells of *Staphylococcus epidermidis* and a MRSA clinical isolate (35, 36). However, although nisin is bactericidal against planktonic cultures, its inhibitory activity is often reduced against sessile cells (27, 37). Therefore, investigating the penetration of antimicrobials through bacterial biofilms could provide valuable information about their therapeutic potential and help to improve antimicrobial dosing regimens (20). Likewise, although previous research has shown that nisin can penetrate bacterial biofilms, these earlier studies could not demonstrate how deep the peptide reached in the biofilm matrix and were not able to differentiate if the effect was bactericidal or bacteriostatic (35, 36).

In the present study, the ability of nisin to penetrate both artificial coverslip biofilms (bacteria entrapped in agarose gel drops attached to a glass slide) and CDC coverslip biofilms of *S. aureus* was monitored by real-time confocal microscopy using a new approach for the preparation and visualization of bacterial biofilms. The sensitivities of planktonic and sessile cells of *S. aureus* to nisin were evaluated by culture-dependent methods and these results were related to the ability of the peptide to modify the permeability of the cytoplasmic membrane of target cells.

## Materials and methods

### Microorganism and growth conditions

*Staphylococcus aureus* AH2547, a strain derivative of *S. aureus* NCTC 8325, was kindly provided by Alexander Horswill, University of Iowa (U.S.A.). *S. aureus* AH2547 contains the plasmid pCM29, which constitutively expresses a green fluorescent protein (GFP) and confers resistance to chloramphenicol. *S. aureus* AH2547 was routinely cultivated overnight at 37 °C in 30 g.l^-1^ tryptic soy broth (TSB) supplemented with chloramphenicol (CHL, Fisher Scientific, Pittsburgh, PA, U.S.A.) at a final concentration of 10 ug.ml^-1^. Incubations were carried out in an orbital shaking incubator at 200 rpm. In order to obtain exponential phase planktonic cultures of *S. aureus* AH2547, a 1 % (v/v) aliquot of an overnight culture (stationary phase planktonic culture) was transferred to TSB media and incubated for 2 h at 37 °C.

### Solutions and antimicrobial agents stock solutions

The sodium phosphate buffer used (PB, 100 μmol.l^-1^, pH 7.0) consisted of an aqueous solution containing, per liter, 4.57 g NaH_2_PO_4_ and 8.78 g Na_2_HPO_4_. Dilution water used for serial culture dilutions consisted of an aqueous solution containing (per liter): 0.405 g MgCl_2_·6H_2_O and 0.0425 g KH_2_PO_4_, with a final pH of 7.2 ± 0.2, prepared according to Method 9050 C.1a (38). Isotonic saline solution used for resuspension of planktonic cells and biofilm cultures consisted of NaCl (0.85 % w/w) prepared in distilled water and autoclaved for 20 min. Nisin stock solutions were prepared from commercial products (2.5 % w/w; Sigma, N5764, Saint Louis, MO, U.S.A.) in sodium phosphate buffer (PB, 100 μmol.l^-1^, pH 7.0) and were stored at 4 °C. A solution of 2-hydroxyethylagarose (4 % w/w; A4018-Sigma-Aldrich, Saint Louis, MO, U.S.A.) was prepared in distilled water, autoclaved for 10 min and stored at room temperature until use. A stock solution of propidium iodide (PI) was resuspended in water at final concentration of 20 μmol.l^-1^. Film Tracer™ Calcein Red-Orange Biofilm Stain (CAM; F10319 Thermo Fisher Scientific, Waltham, MA, U.S.A.) was resuspended in DMSO at final concentration of 0.416 ug.ml^-1^. Both dye solutions were stored at −20 °C until use.

### Minimum inhibitory concentration

Minimum inhibitory concentrations (MIC) for nisin were determined using the CLSI micro-broth dilution method (39). *S. aureus* AH2547 was grown as previously described. The nisin concentration range tested varied from 0.1 to 12.5 μmol.l^-1^. TSB media was used and the inoculum was standardized to 10^5^ CFU.ml^-1^. The MIC was defined as the lowest concentration of nisin that caused complete growth inhibition of after 48 h of incubation at 37°C. Results represent the mean values ± standard deviation (SD) of two independent determinations performed with three replicates.

### Calcein-AM staining

Overnight planktonic cultures of *S. aureus* AH2547 were centrifuged and resuspended in 1 ml of sterile water. These cell suspensions were stained with 20 μl of CAM stock solution for one hour at room temperature. Excess CAM was removed by centrifugation. Similarly, CDC coverslip biofilms were gently immersed into 2.5 ml PB added with 50 μl of CAM stock solution for one hour at room temperature. Excess CAM was removed by rinsing with PB without CAM.

### Nisin effect on cytoplasmic calcein retention

Overnight planktonic cultures of *S. aureus* AH2547 were stained with CAM as described above. The stained cultures were exposed to 15 or 150 μmol.l^-1^ nisin for 60 min at room temperature. The control treatment was performed using PB without nisin. After the treatments, the samples were filtered (Lab System 3 – Millipore Corporation, Bedford, MA, U.S.A.) and immobilized onto polycarbonate black membranes (25 mm, 0.22 μM pore size, Osmonics INC, Minnetonka, MN) and transferred to glass slides (Fisherbrand Superfrost/Plus microscope slides, Fisher Scientific, Pittsburgh, PA, U.S.A.). Samples were examined microscopically (more details below). The experiment was performed with three biological replicates.

### Artificial coverslip biofilm

CAM-stained overnight planktonic cultures of *S. aureus* AH2547 were transferred to a 4 % solution of 2-hydroxyethylagarose (A4018-Sigma-Aldrich, St. Louis, MO, U.S.A.) at 50 ºC ± 5 ºC with a 1:1 proportion. The mixture was immediately vortexed and small drops were transferred onto circular glass coverslips to create the artificial coverslip biofilms (1.5 Micro Cover Glass-72225-01; Electron Microscopy Sciences, Hatfield, PA, U.S.A.). Syringes (1 ml) with 25 G 1½ needles (305127-Precisionglide) were used to dispense the drops. The glass coverslips with artificial biofilm drops were covered with PB until use. The experiment was performed with three biological replicates.

### CDC biofilms

Sessile *S. aureus* AH2547 cultures were grown in CDC biofilm reactors (Biosurface Technologies Inc., Bozeman, MT, U.S.A.) (40) on glass coupons. Briefly, 1 ml of an overnight planktonic culture was transferred to a reactor containing 400 ml of full-strength TSB media (30 g.l^-1^) and 10 ug.ml^-1^ of CHL (Fisher Scientific, Pittsburgh, PA, U.S.A.). Cultures were incubated at 37 °C with constant stirring (125 rpm). After 24 h of batch growth, the continuous flow (11.7 ml.min^-1^) of one-tenth-strength TSB (3 g.l^-1^) was started and continued for an additional 48 h. The CDC coupons were removed for analysis after 48 h of continuous flow. The outside surface (low shear side) of CDC coupons was used in biofilm experiments. When appropriate, glass coverslips (1.5 Micro Cover Glass-72225-01; Electron Microscopy Sciences, Hatfield, PA, USA) were attached to the outside of CDC - polypropylene coupon holders supported by an glass microscope slide with latex elastic bands (Intra-oral Latex Elastics) and then, at the end of incubation, transferred to the flow cell reactor in order to evaluate nisin penetration as shown diagrammatically in Fig. 1.

**Figure 1.**
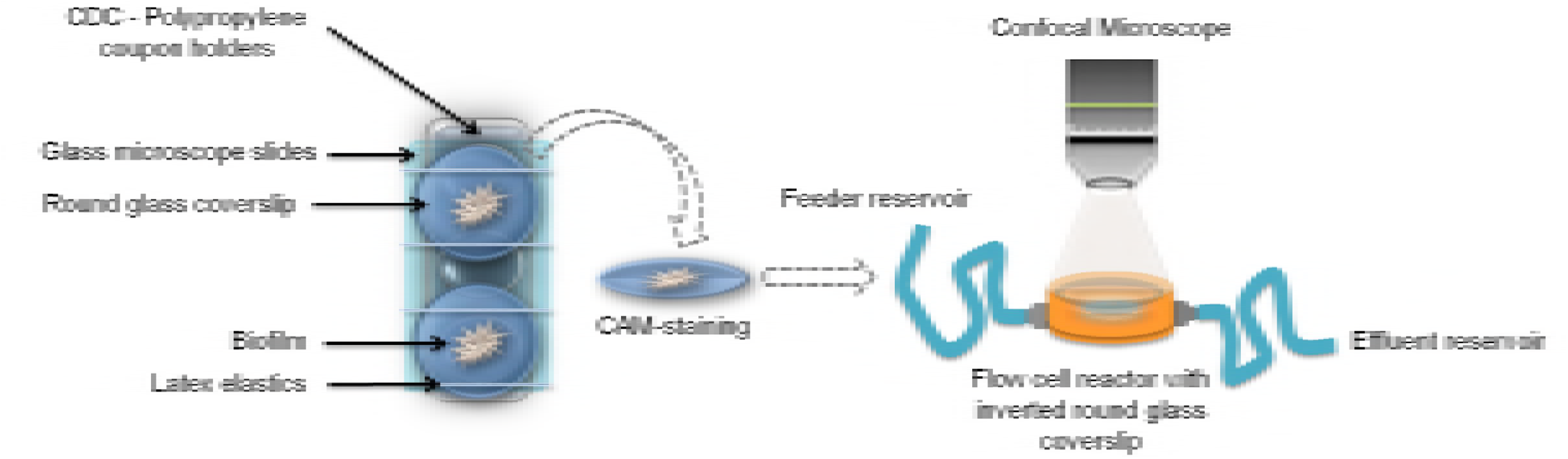
Experimental procedures to obtain and treat CDC coverslip biofilms with nisin. *S. aureus* was grown in a CDC reactor containing round glass coverslips fastened to microscope slides with rubber bands. The CDC coverslip biofilms were stained with CAM and the inverted glass coverslip containing the biofilm was placed in a flow cell reactor. Nisin at 15 μmol.l^-1^ was supplied to the flow cell at a rate of 2 ml.min^-1^ during 1 h at room temperature and the fluorescence at the bottom of the biofilm was monitored over time with a confocal microscope.

### Flow cell reactor treatment and TL-CLSM analysis

A flow cell reactor fitted to receive as part of its cover a glass coverslip was placed on the stand of a confocal scanning laser microscope (CLSM) and connected to a peristaltic pump by silicone tubing. One end of the first tubing section was connected to the feeder reservoir and the other end of the second tubing section was connected to an effluent reservoir as shown in Fig 1. Glass coverslips containing artificial biofilms or CDC biofilms were previously stained with calcein dye and then gently placed, inverted, into the flow cell reactor, forming the cover. After the remainder of the cover was screwed on, the internal space of the flow cell reactor was carefully filled with PB, to avoid bubble formation. The outside face of the glass coverslip was gently cleaned and the focus plane on the CLSM was adjusted. Next, a solution of 15 μmol.l^-1^ nisin was pumped through the flow cell at a rate of 2 ml per min for 1 h at room temperature. PB without nisin addition was used as the control treatment. Throughout the experiments, the GFP and calcein fluorescence were monitored using bright field and fluorescence imaging (TL-CSLM) on the confocal microscope, where six frames were recorded per second. The experiment was performed with three biological replicates.

### PI staining and cryosectioning of CDC biofilms

Upon treatment completion, biofilms on the outside surfaces of CDC coupons were rinsed and stained with propidium iodide (PI) as follows: 150 μl of stock PI solution were pipetted directly onto the biofilm surface and the coupons were allowed to sit for 10 min at room temperature. Next, excess PI was gently removed by rinsing with sterile water and samples were cryoembedded. Tissue-Tek O.C.T. compound (Miles Laboratories, Elkhart, Indiana, U.S.A.) was added on the top of the coupons before these being placed on a slab of dry ice and frozen. The frozen O.C.T. - embedded biofilm was removed from the coupon surface, and re-embedded (41). The embedded samples were stored at −80 ºC until use. Samples were cryosectioned in 5 μm slices on a Leica CM1800 cryostat and transferred to Superfrost Plus microscope slides (Fisher Scientific, Pittsburgh, PA, U.S.A.) and stored at −80 ºC until microscopic analysis (more details below). Samples without PI staining were used as controls. The experiment was performed with two biological replicates and three technical replicates.

### Microscopy analyses

Epi-fluorescent images were captured on a Nikon Eclipse E-800 microscope using a Photometrics^®^ CoolSNAP MYO CCD camera and MetaVue software (Universal Imaging Co., Downingtown, PA, U.S.A.). Standard FITC and TRITC fluorescence filter cubes were used to visualize the GFP fluorescence (green fluorescence) and PI/Calcein fluorescence (red fluorescence), respectively, with a 20× 0.75NA dry objective. Confocal images were acquired using a Leica TCS-SP5 confocal scanning laser microscope (CSLM), with 10× objective lens. The 488 nm and 561 lasers were used for excitation of fluorophores, and the detector bandwidth was set to collect GFP and calcein emission fluorescence between 500-550 nm and 575-675 nm, respectively. The fluorescence intensity was measured using Metamorph and Imaris software (Bitplane Scientific Software, Saint Paul, MN, U.S.A.). Twelve regions measuring 10 μm per side each were randomly distributed in the center, middle and edge zone of *S. aureus* biofilm fields of view. A single plane of focus for each sample was selected and the GFP and calcein fluorescence intensity were measured for all twelve regions during the entire treatment period. In order to allow pooling of background-corrected data from different experiments, four additional regions were selected in areas without biofilm and the mean background fluorescence intensity was subtracted from the measured of GFP and calcein values. The fluorescence values were normalized as the relative intensity by dividing the values of GFP intensity by values of calcein intensity at that specific location.

### Nisin susceptibility of homogeneous cultures

Nisin susceptibilities of exponential and stationary phase planktonic and sessile *S. aureus* cultures were compared. Planktonic growth conditions were performed as described above and the sessile cultures were obtained from the CDC biofilm. Planktonic cultures were centrifuged (2500 × *g*, 8 min at 4 ºC), resuspended in isotonic saline solution and reserved at room temperature until use. The sessile cells were detached from the CDC coupon using a wooden scraper (see below in the viable cell count of CDC biofilm subheading) and resuspended in isotonic saline solution. Both planktonic and sessile cultures were homogenized for 30 sec (5,000 rpm; IKA T25 digital Ultra Turrax) at room temperature and the optical densities at 600 nm were standardized to 0.05. Aliquots of 1 ml from each culture were distributed in centrifuge microtubes and treated with 15 μmol.l^-1^ of nisin for 60 min at room temperature. PB without nisin was used as the control. Samples were removed at 0, 5, 15, 30 and 60 min after exposure to nisin. Additionally, cells were treated for up to 60 min with 150 μmol.l^-1^ of nisin. Samples were then washed by centrifugation (2500 × *g*, 8 min at 4 ºC) to remove the antimicrobial and the cell pellets were resuspended in isotonic saline solution. In order to enumerate the viable cells, serial dilutions were performed and 20 μl from each dilution were transferred onto TSA plates followed by incubation at 37 ºC (42). Plates showing colonies after 24 h of incubation were enumerated and the number of colony forming units (CFU) was calculated. The experiment was performed with two biological replicates and two technical replicates.

### Nisin susceptibility of CDC biofilm

*S. aureus* AH2547 biofilms were treated at room temperature with 150 μl of 15 μmol.l^-1^ nisin, which covered the entire upper surface of the CDC coupon. PB without nisin was used as the control treatment. Samples were removed after 0, 5, 15, 30 and 60 min of nisin treatment, rinsed with the same volume of sterile distilled water and the enumeration of viable cells on the CDC biofilm was performed as described below. When appropriated, biofilm samples were submitted to cryosection analyses as described above. This experiment was performed with two biological replicates and two technical replicates.

### Viable cell count of CDC biofilms

In order to determine viable cells of *S. aureus* in CDC biofilms, sessile cells were enumerated as previously described (40), with the modifications described below. Biofilm from the outward-facing side of the CDC coupon were scraped with a sterile wooden applicator stick. On average, four applicator sticks were used for each coupon. Each stick was rinsed and stirred into the same 10 ml volume of isotonic saline solution. Solutions were homogenized for 30 sec (5,000 rpm; IKA T25 digital Ultra Turrax) and serial dilutions were performed. Aliquots of 20 μl from each dilution were transferred onto TSA plates and colonies that appeared after 24 h of incubation at 37 ºC were counted (42).

### Statistical analysis

Statistical analysis was performed using GraphPad Prism (GraphPad Prism Software Inc., San Diego, CA). Data from nisin susceptibility tests were presented as the mean values ± SD. Significance of interaction effects between nisin and time of treatment was determined using a two-way repeated-measures ANOVA test. Post-hoc pairwise multiple comparison procedures included the Bonferroni test (comparing the mean of a treated group) and Student-Newman-Keuls test (comparing the factor level mean within a treated group). The probability level for statistical significance was *P*<0.05.

## Results

The MIC of nisin against *S. aureus* AH2547 was 1.56 μmol.l^-1^ (data not shown). To account for the physiological heterogeneity between planktonic and sessile cells and the increased resistance to antimicrobials of biofilm-grown cells, subsequent killing assays were performed using 15 μmol.l^-1^ (~10× MIC) or 150 μmol.l^-1^ nisin (100× MIC). To verify the ability of nisin to cause membrane permeabilization of the target cells with consequent loss of calcein (a fluorescent intracellular dye), overnight planktonic cultures of *S. aureus* AH2547 stained with CAM were exposed to nisin. First, the absence of red fluorescence was confirmed in a negative control treatment performed with cultures of *S. aureus* AH2547 not stained with CAM but treated with nisin (Fig. 2A). These cultures fully retained their GFP fluorescence (green fluorescence) after nisin treatment, confirming that nisin does not cause significant loss of GFP fluorescence during 60 min of treatment. Additionally, cultures of *S. aureus* AH2547 stained with CAM and treated with buffer without nisin retained most of their red fluorescence (Fig. 2B). However, when *S. aureus* AH2547 cells stained with CAM were treated with nisin (15 μmol.l^-1^ nisin), an almost complete loss of intracellular calcein was observed (Fig. 2C). This effect was even more dramatic when *S. aureus* cells were exposed to 150 μmol.l^-1^ nisin (Fig. 2D). These results confirmed that calcein could be a useful marker to monitor the antimicrobial activity of nisin on *S. aureus* AH2547 cultures.

**Figure 2.**
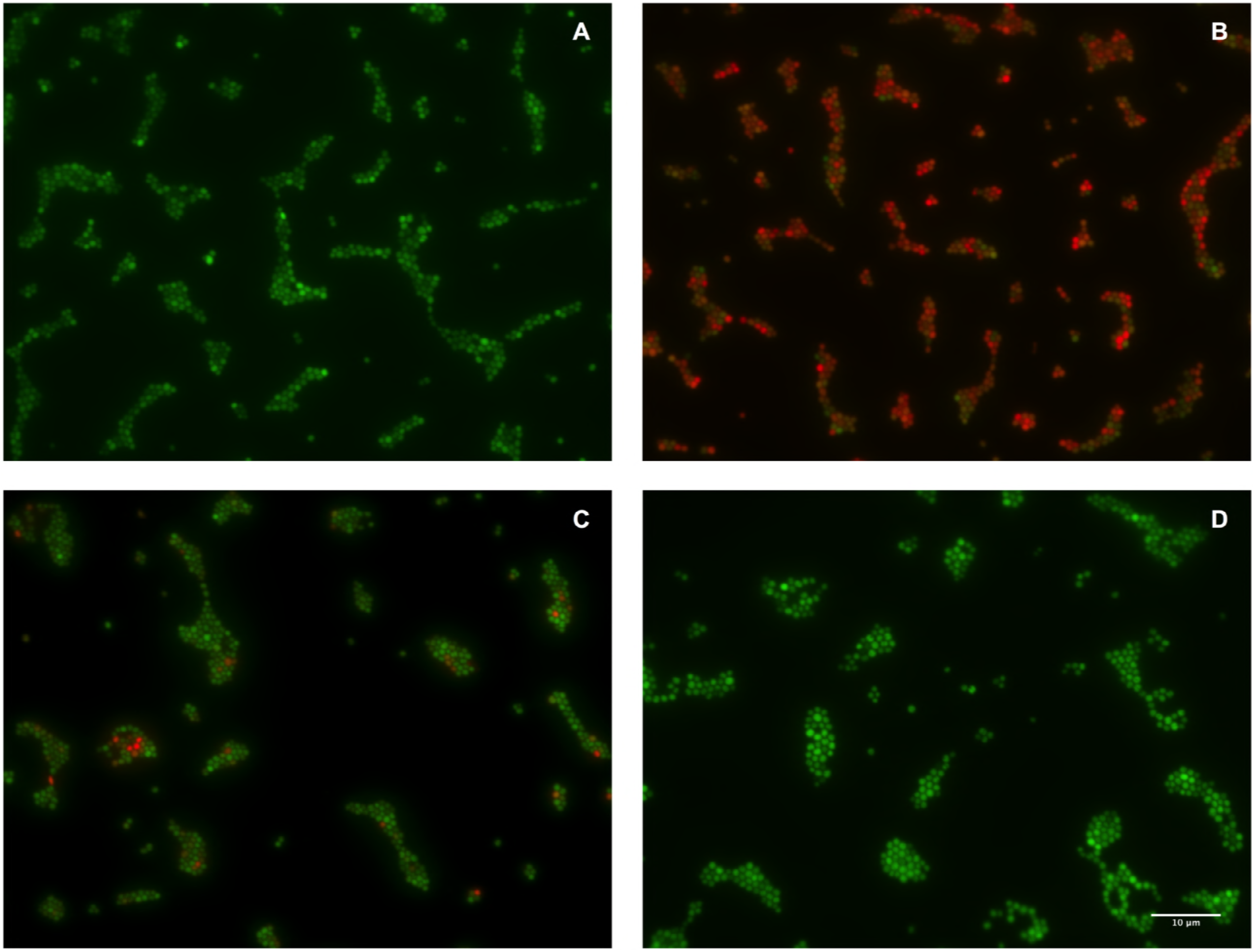
Effect of nisin against calcein-AM (CAM) stained cultures of *S. aureus* AH2547. Cell suspensions containing approximately 10^5^ CFU.ml^-1^ of overnight planktonic *S. aureus* AH2547 were stained with calcein-AM and incubated at room temperature with nisin (15 μmol.l^-1^) for 60 min. The cultures were immobilized onto black polycarbonate membranes and visualized by fluorescence microscopy (100× magnification). A – Culture not stained with CAM and exposed to nisin; B – Culture stained with CAM and not exposed to nisin; C – Culture stained with CAM and treated with 15 μmol.l^-1^ nisin; D – Culture stained with CAM and treated with 150 μmol.l^-1^ nisin. Three independent evaluations were performed for each treatment and the images are representative of the observations fields for each treatment.

To adjust experimental conditions and to monitor the activity of nisin with consequent loss of intracellular CAM in immobilized cells under a flow treatment, we developed an experimental approach to allow direct real-time evaluation of the bacterial biofilms treated with nisin. Stationary phase cultures were stained with CAM and immobilized in artificial coverslip biofilms made of agarose gels in a custom-made flow cell and fluorescence was monitored with TL-CLSM. (Fig. 3). Both calcein and GFP fluorescence in untreated artificial coverslip biofilms were consistently retained throughout the treatment confirming minimal photobleaching and photodamage over treatment time (Fig. 3A-F, Movie S1). However, when artificial coverslip biofilms were treated with 15 μmol.l^-1^ nisin, the red calcein fluorescence was progressively lost while the green GFP fluorescence was maintained over time, indicating extensive membrane permeabilization (Fig. 3G-M, Movie S2). Red fluorescence dissipated uniformly at the edges and at the center of the artificial biofilm.

**Figure 3.**
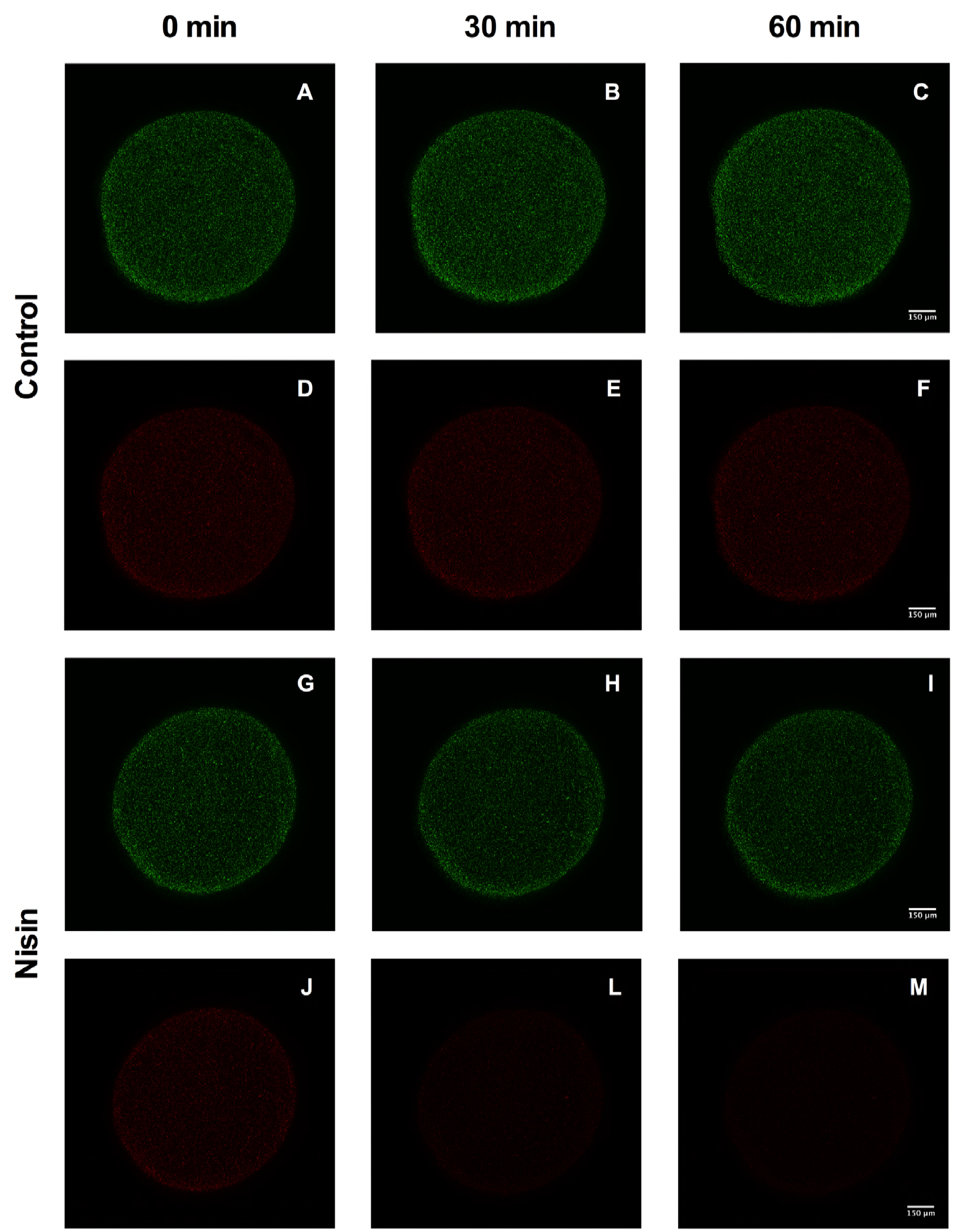
Nisin flow cell treatment of *S. aureus* AH2547 artificial coverslip biofilms. Sequential images from GFP fluorescence (A-C, G-I) and calcein fluorescence (D-F, J-M) captured by TL-CSLM of *S. aureus* artificial coverslip biofilms under flow cell treatment in a rate of 2 ml.min^-1^ at room temperature at 0, 30 and 60 min. Control treatments (C) were performed with phosphate buffer without nisin (A-F) and nisin treatment (N) was performed with phosphate buffer with 15 μmol.l^-1^ nisin (G-M). The focal plane in these experiments is located near the bottom of the artificial coverslip biofilms (the side close to the glass). The experiment was performed with three biological replicates and the images are representative for each treatment.

*S. aureus* AH2547 CDC coverslip biofilms were then submitted to flow cell reactor treatments similar to what was applied to the artificial coverslip (Fig. 4). In agreement with the previous assay, GFP fluorescence was retained in both control and nisin treated specimens over the incubation time. Although the red calcein fluorescence varied over incubation time in both control and nisin treatments, nisin-treated biofilms lost the red calcein fluorescence much faster and thoroughly than the control treatment (Fig. 4D-F, Movie S3 and S4). These observations confirmed that nisin could penetrate into the biofilm and disrupt the integrity of the cytoplasmic membrane of *S. aureus* sessile cells. Membrane permeabilization in both artificial biofilms and CDC coverslip biofilms in the flow cell reactors were quantified by the change in red (calcein) to green (GFP) fluorescence ratio (Fig. 5). In the artificial biofilm assay, the decrease in red to green fluorescence ratio was much greater in the nisin treated samples (85 % reduction after 60 min exposure) than in untreated controls (15 % reduction after 60 min exposure). The time for the red to green fluorescence ratio to drop to half of its initial value in artificial *S. aureus* biofilm exposed to nisin was 28 min (Fig. 5). In the CDC coverslip biofilm assay, control treatments showed a gradual reduction of the fluorescence ratio during continuous flow treatment with approximately 17 % loss in the red to green fluorescence ratio after 15 min. In contrast, the CDC coverslip biofilm treated with nisin showed a more pronounced decrease in fluorescence ratio with approximately 75 % of the red to green fluorescence ratio over the same 15 min interval. The time required to reach half of the initial value of the red to green fluorescence ratio in *S. aureus* CDC coverslip biofilms was 11 min when exposed to nisin (Fig. 5B).

**Figure 4.**
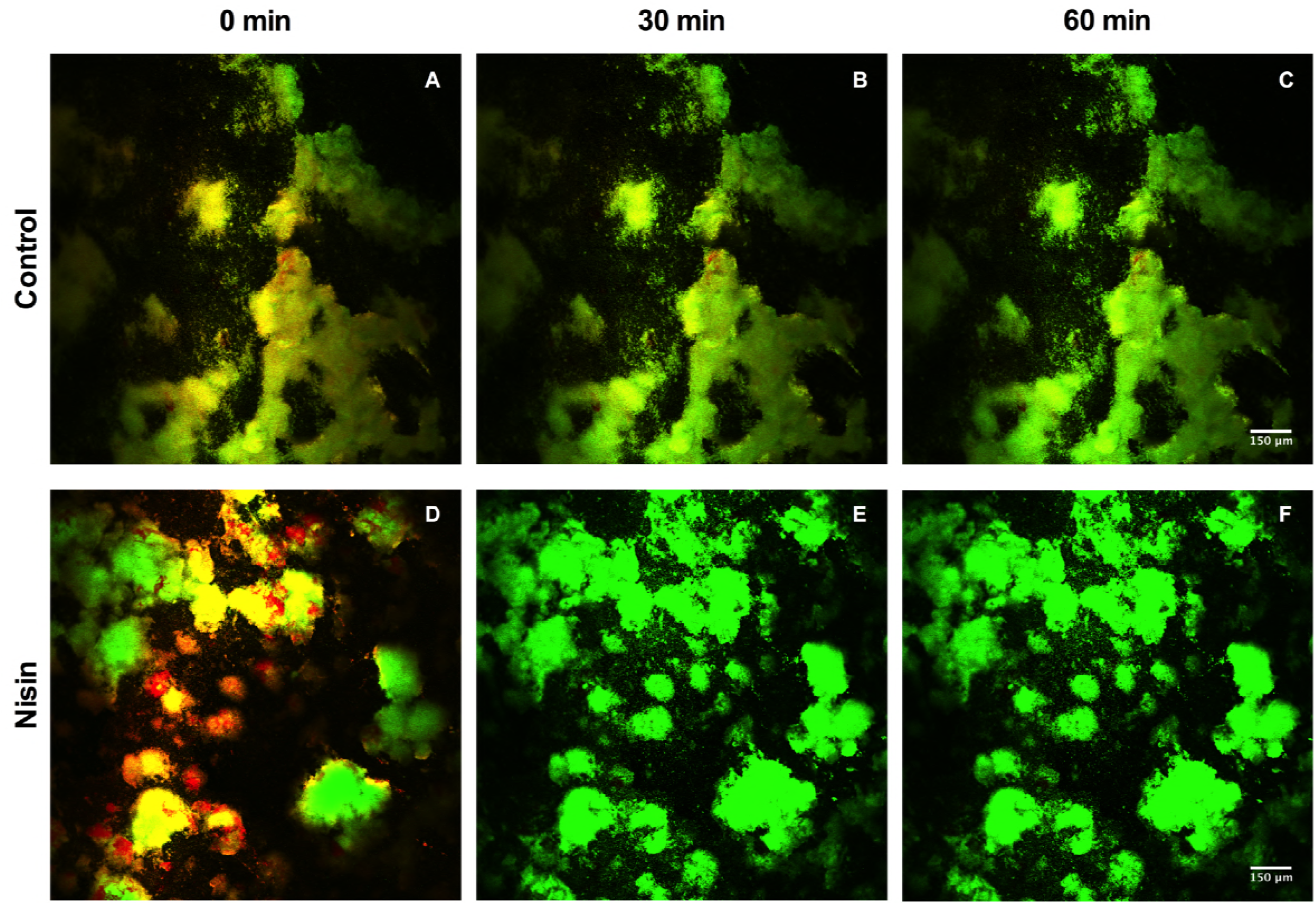
Nisin flow cell treatment of *S. aureus* AH2547 CDC coverslip biofilms. Sequential images of *S. aureus* CDC coverslip biofilms under flow cell treatment in a rate of 2 ml.min^-1^ at room temperature at 0, 30 an 60 min. Control treatments (C) were performed with phosphate buffer without nisin (A-C) and nisin treatment were performed with phosphate buffer with 15 μmol.l^-1^ nisin (D-F). The GFP (green) and calcein (red) fluorescence were captured by TL-CSLM assay and the images were superimposed. The focal plane in these experiments is located near the bottom of the biofilm. Three independent evaluations were performed for each treatment and the selected images are representative for each treatment.

**Figure 5.**
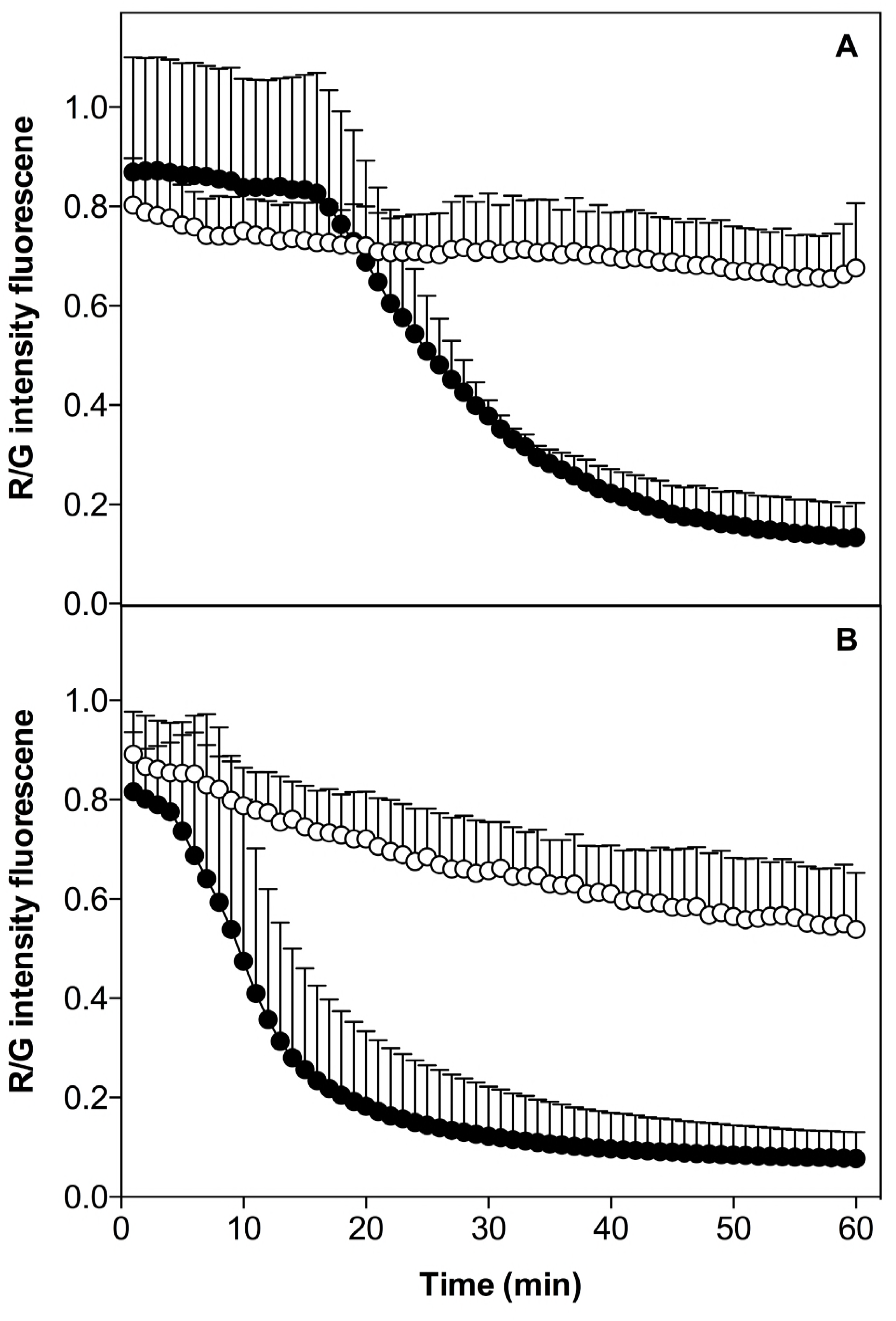
Calcein to GFP fluorescence ratio from *S. aureus* biofilms under flow cell reactor treatment. Calcein and GFP fluorescence of *S. aureus* AH2547 biofilms under flow cell treatment in a rate of 2 ml.min^-1^ during 1 h at room temperature were detected by CSLM and quantified using Metamorph software. Artificial coverslip biofilms (A) and CDC coverslip biofilms (B) of *S. aureus* AH2547 is shown. Control treatments (white circles) were performed with phosphate buffer without nisin and nisin treatment (black circles) were performed with phosphate buffer with 15 μmol.l^-1^ nisin. Fluorescence intensities were measured at twelve locations distributed randomly among the center, middle and edge zone of the biofilm. Data represent the means and standard deviations of three replicates.

To further investigate the ability of nisin to penetrate bacterial biofilms and affect cell viability, biofilms of *S. aureus* AH2547 were grown on CDC coupons in the CDC biofilm reactor, treated with nisin, and stained with PI. Biofilm cryosection samples were analyzed by confocal microscopy (Fig. 6). Cells with damaged membranes are expected to take in the red PI whereas intact cells should not stain red. Expected morphological variations were observed among the CDC *S. aureus* biofilm specimens, such as differences in biofilm thickness and the presence or absence of robust biofilm clumps. Absence of red fluorescence was confirmed on control samples without nisin or PI staining (Fig. 6A). Additionally, as described above for planktonic cells, nisin treatment did not change the GFP fluorescence in biofilm cells (Fig. 6B). Untreated biofilm samples stained with PI, showed some red fluorescence even though the cells had not been exposed to nisin (Fig. 6C). These results suggested that some bacteria in the biofilm were already dead or were naturally more permeable to PI. In contrast, most biofilm samples treated with nisin were completely stained by PI (Fig. 6D). Nonetheless, it should be pointed out that, in only a few cases, differences in biofilm thickness or the presence of clumps in the biofilm affected PI staining of nisin-treated samples (Fig. 6 E).

**Figure 6.**
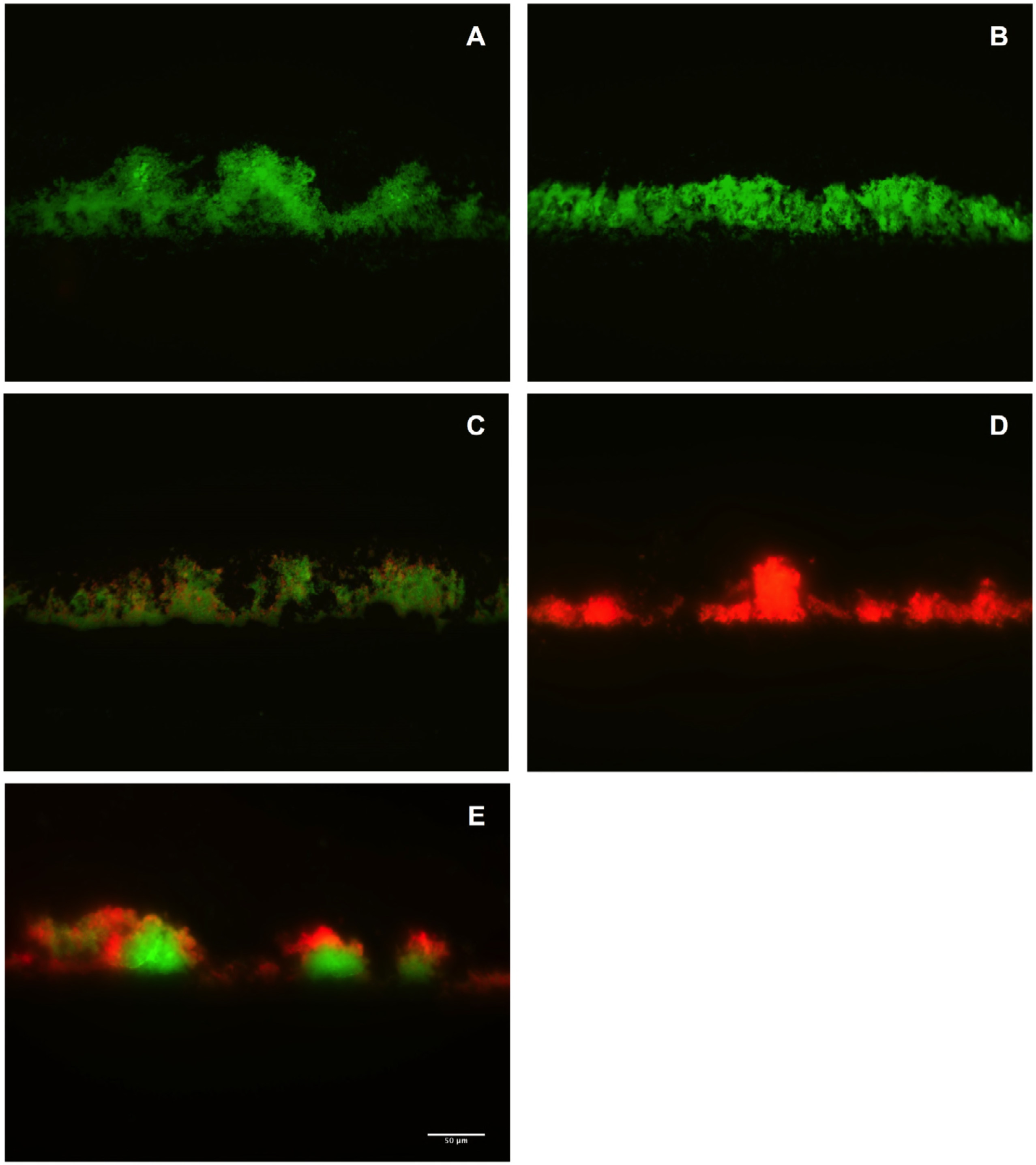
Fluorescence microscopy images of *S. aureus* AH2547 CDC biofilms exposed to nisin. Before cryosectioning, *S. aureus* AH2547 CDC biofilms expressing GFP were treated with 15 μmol.l^-1^ nisin during 1 h at room temperature and stained with PI. Cryosectioned biofilm samples were immobilized onto microscope slides and visualized by fluorescence microscopy (20× magnification). The control treatments were performed with phosphate buffer without nisin: A – Samples not stained with PI and not exposed to nisin (control 1), B – Samples not stained with PI and exposed to nisin (control 2). C – Samples stained with PI and not exposed to nisin (control 3), Panels D and E show the *S. aureus* biofilms that were stained with PI and treated with nisin. This experiment was performed with two biological replicates and the images are representative fields for each treatment.

The viabilities of *S. aureus* AH2547 cultures in different physiological states were particularly affected when exposed to different nisin concentrations (15 μmol.l^-1^ and 150 μmol.l^-1^). Relevant changes in viable cell number could be observed at different time intervals for most samples, demonstrating significant interaction between nisin and exposure time (*P*<0.001) (Fig. 7). In general, planktonic cultures showed high susceptibility when compared to sessile cells. Homogenized exponential planktonic cells (HEP) were highly sensitive to 15 μmol.l^-1^ of nisin after only 5 min of exposure. HEP viability decreased below the detection limit with a log reduction of at least 6.99 ± 0.78 log CFU.ml^-1^. As expected, a similar decrease in viability was observed when a 10-fold higher nisin concentration was applied to HEP cells (*P*<0.001). Homogenized stationary planktonic cells (HSP) were much less sensitive to nisin than HEP (*P*<0.001). The maximum log reduction in HSP viability upon exposure to 15 μmol.l^-1^ nisin was 1.64 ± 0.50 CFU.ml^-1^. The increase in nisin concentration (15 to 150 μmol.l^-1^) caused greater bactericidal activity (*P*<0.001) against HSP cells, with a log reduction of 3.6 ± 0.40 CFU.ml^-1^. In contrast to the planktonic cultures, biofilm cultures were less susceptible to nisin than homogenized planktonic cultures. Maximum reduction in bacterial viability of homogenized biofilms (HB) and intact biofilms (IB) were always below 2 log cycles, even when the cultures were exposed to a very high concentration of nisin (150 μmol.l^-1^).

**Figure 7.**
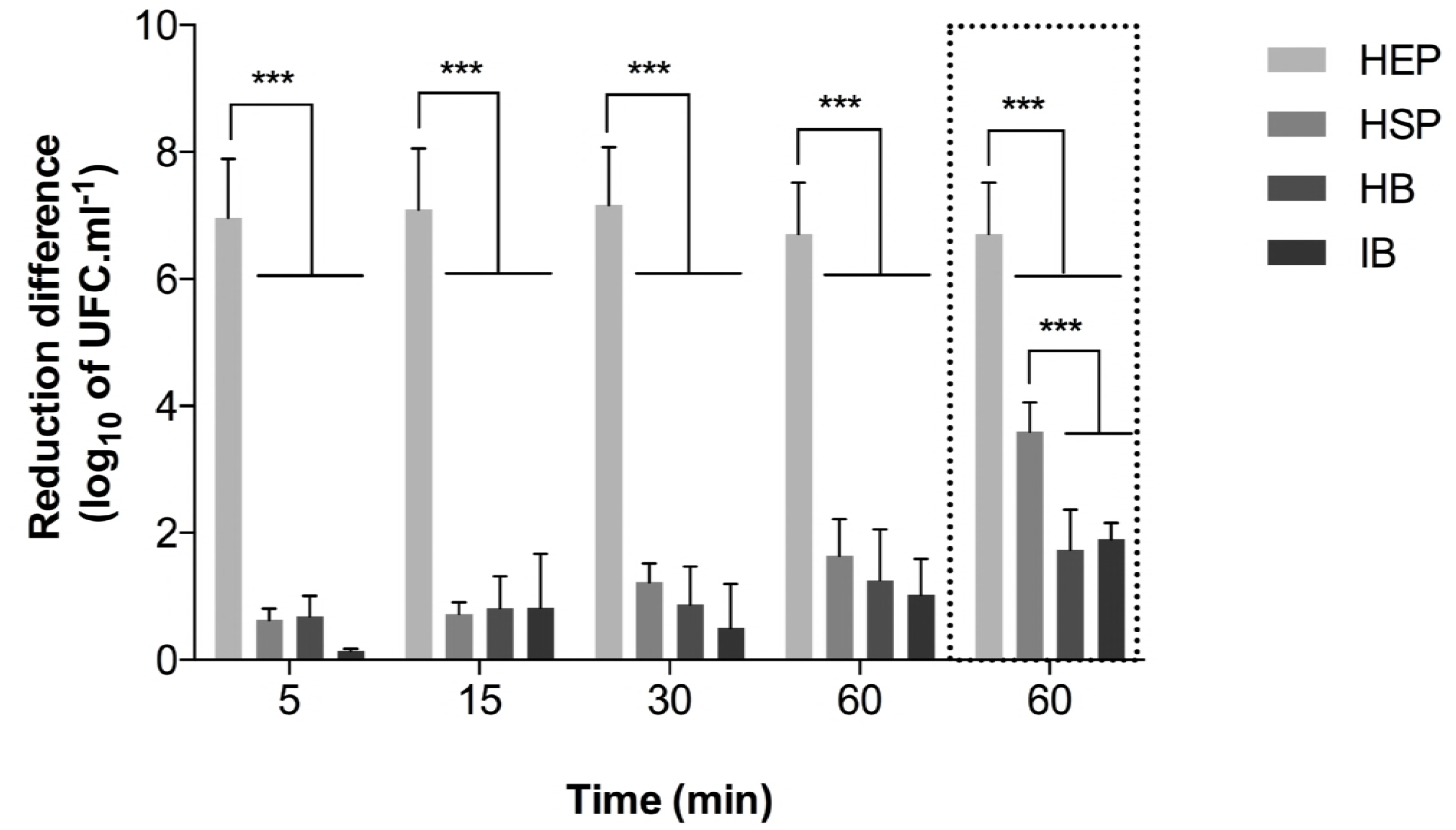
Susceptibility of planktonic and sessile cells of *Staphylococcus aureus* AH2547 to nisin. Nisin (15 μmol.l^-1^) was incubated with homogenized exponential planktonic (HEP) cells, homogenized stationary planktonic (HSP) cells, homogenized CDC biofilm (HB) and intact CDC biofilm (IB). Viabilities were determined by enumeration of CFU.ml^-1^ following different incubation times (0, 5, 15, 30 and 60 min). When indicated, cultures were treated with 150 μmol.l^-1^ during 60 min of incubation (bars delimited by a dotted line). Control treatments were performed with phosphate buffer without nisin. The data represents the difference in viable cell number between the control and treated samples. Mean values and the standard deviations of triplicate independent determinations are shown. Viability detection limit corresponded to 4 × 10^2^ CFU.ml^-1^. Horizontal lines above plot indicate significant differences and asterisks indicate a level of significance (\*\**P*<0.001).

## Discussion

In this work, we experimentally confirmed that nisin, a polycyclic lantibiotic widely recognized as an effective antimicrobial agent against several bacterial pathogens, could penetrate into a bacterial biofilm and cause membrane permeabilization of *S. aureus* cells. We also showed that membrane integrity loss correlated with a decrease in bacterial viability. To achieve this, we used a flow cell reactor coupled with real-time TL-CLSM to provide a direct evaluation of nisin penetration through *S. aureus* biofilms. Calcein release was used to monitor the ability of nisin to form pores in the cytoplasmic membranes of *S. aureus* cells (Fig. 2). Calcein dye shows a stronger fluorescence under cytoplasmic conditions, but a loss in cell membrane integrity causes a rapid release of the dye (43, 44). Previous work demonstrated that calcein can easily diffuse through bacterial biofilms without disrupting their structure, which was an useful feature for the present study (44). In addition, to overcome the limitations of light penetration through the entire biofilm while it was being exposed to nisin, we changed the location of the focal plane at the bottom of the biofilm and near the coverslip interface. This allowed us to monitor the GFP and calcein fluorescence in the biofilm, confirming that nisin was able to fully penetrate established biofilms and kill sessile cells. Therefore, in our flow cells, the deepest regions of the biofilm (i.e. the bottom of the structure), was at the top of the flow cell, against the inner side of the coverslip, which allowed us to effectively image the penetration of the antimicrobial peptide into the bacterial biofilm. These results also indicated that nisin did not mechanically disrupt the artificial or the CDC biofilms (Fig. 3 and 4), corroborating with previous observations (35, 36, 44).

Our artificial biofilms containing bacteria entrapped into agarose gels were much thicker than the CDC coverslip biofilms, a feature that appeared to affect the release of calcein from the biofilm. In the artificial biofilm, calcein loss was not significant for at least 17 min after the start of the nisin treatment compared to the CDC coverslip biofilm, indicating that the effect of the antimicrobial peptide on the biofilm was being influenced by microbial biomass and the thickness of the extracellular matrix (17). It is also important to note that some decrease in calcein fluorescence was observed in control treatments, mainly in the CDC coverslip biofilms, which might be caused by active esterases released into the extracellular matrix (from cell lysis) that could hydrolyse the calcein in the biofilm matrix followed by its subsequent loss (wash out) of the fluorescent dye due to the flow treatment (45, 46)

PI staining confirmed that nisin could penetrate the bacterial biofilm and affect the membrane permeability of sessile *S. aureus* cells (Fig. 6). PI is an intercalating agent that shows stronger red fluorescence when bound to DNA and its diffusion into the cells only occurs if the cytoplasmic membrane becomes permeable, which makes this molecule a suitable marker to assess cell viability loss in *in vitro* killing assays (47, 48). Qualitative analysis of the cells stained with PI by fluorescence microscopy demonstrated that nisin-treated *S. aureus* biofilms were quickly stained by PI. Nonetheless, some variability of PI staining was observed when biofilm clumps were present, indicating that biofilm biomass/morphology can affect the efficacy of nisin diffusion through the extracellular matrix.

Enumeration of viable *S. aureus* cells with and without nisin treatment under the same conditions applied for the PI staining was used to evaluate the viability of *S. aureus* cells in established biofilms (CDC reactor). Approximately 83.3 % of the sessile *S. aureus* cells were killed when intact biofilm samples were treated with 15 μmol.l^-1^ nisin, while a decrease of approximately 98.6% of the viable cell number occurred when the nisin concentration was 150 μmol.l^-1^. These latter results also indicated that cell physiology (exponential versus stationary phase) has a significant impact on nisin susceptibility in *S. aureus* cultures. Taken together, these results agree with previous observations that demonstrated the differences in antimicrobial susceptibility between exponential versus stationary cells in different species of bacteria (37, 49–51).

In this study, nisin was bactericidal against exponentially growing *S. aureus* cultures and against sessile (biofilm-grown) and stationary cell cultures (Fig. 7). However, the magnitude of these effects were distinct, with exponential cells showing a much greater reduction (≥ 7 log cycles) in viable cell numbers compared to sessile and stationary cells (~ 2 log cycle). Biofilm cells shows great physiological heterogeneity, but adaptive responses that resembles what occurs during the stationary phase is expected to predominate in the attached microorganisms, with most cells being characterized by a slow growth rate and a low cell-wall turnover rate (9, 52). In this scenario, although nisin could make pores in the cytoplasmic membrane of *S. aureus* sessile cells, protective stress responses could limit the efficacy of the antimicrobial peptide and prevent inhibition of cell wall biosynthesis in biofilm cells.

Moreover, although in rod-shaped bacteria new peptidoglycan precursors can be added either into their lateral wall or at their division septum, in *S. aureus* the cell wall biosynthesis occurs predominantly in the division septum, which is rich in lipid II (53, 54). Lipid II is the main target of nisin in bacterial cells that act as an anchor during the pore formation in the cytoplasmic membrane (34, 51, 55). This uneven distribution of lipid II during cell wall biosynthesis in *S. aureus* was also confirmed with telavancin, a semisynthetic derivative of vancomycin with high affinity to lipid II, which was shown to bind mostly to lipid II in the division septum of exponentially growing *S. aureus* cells causing membrane depolarization (54).

Lipid II is essential for the transport of peptidoglycan monomers during cell wall biosynthesis and serve as a membrane anchor to nisin during pore formation on sensitive organisms (56, 57). The N-terminal lanthionine rings of nisin interact with the pyrophosphate group of lipid II via direct hydrogen bonding allowing the C-terminal region of the peptide to insert into the cytoplasmic membrane (56). When nisin-lipid II complexes are formed in the cytoplasmic membrane, a dramatic efflux of cytoplasmic compounds is induced resulting in membrane depolarization with consequent cellular death (58). In addition to the ability of nisin to form large and stable pores (diameters varying from 2 to 2.5 nm) in the cytoplasmic membrane, specific interactions between nisin and lipid II prevent the incorporation of newly synthesized glycan chains into the existing sacculus thus inhibiting cell wall biosynthesis (56, 59).

Since direct evaluation confirmed that nisin was able to penetrate *S. aureus* biofilms and damage *S. aureus* sessile cells, the variation in nisin susceptibility appeared to be mainly related to phenotypic (physiological) differences between the sessile and planktonic cultures. Therefore, strategies such as the use of higher nisin concentrations or its association with other antimicrobial agents could improve the bactericidal activity against *S. aureus* biofilms. In this study, when the concentration of the antimicrobial peptide was increased, a significant effect (*P*<0.05) in viability loss of intact biofilm cells was observed compared to the lower dose (Fig. 7).

In conclusion, we provide real time evidence through direct microscopic visualization that nisin is able to penetrate into *S. aureus* biofilms and kill sessile cells of a relevant human and animal pathogen. Considering the similarities in sensitivity of HSP and HB cells to nisin, the relevance of using stationary phase cultures in susceptibility tests to get a better estimation of the effects of antimicrobial agents against bacterial biofilms should be emphasized, specially against multidrug resistant bacteria. Based on evidence that biofilms show a wide array of resistance mechanisms, our new approach to assess biofilm susceptibility could improve the evaluation of susceptibility breakpoints of promising antimicrobials. Over the years, a combination of traditional techniques and confocal microscopy has promoted significant advances in the field and further investigations using the flow cell reactor could provide more realistic assessments of the therapeutic effects of antimicrobial agents against sessile cells of microbial pathogens.

## Acknowledgements

This work was supported by the Fundação de Amparo a Pesquisa do Estado de Minas Gerais (FAPEMIG; Belo Horizonte, Brazil) and Coordenação de Aperfeiçoamento de Pessoal de Nível Superior (CAPES; Brasília, Brazil). F.G.S. received a fellowship from the Conselho Nacional de Desenvolvimento Científico e Tecnológico (CNPq; Brasília, Brazil). Microscopy at the Center for Biofilm Engineering was made possible by awards from the Murdock Charitable Trust and the National Science Foundation (CBET 1039785).

